## Supplementary Material for "Prior dengue immunity enhances Zika virus infection of the maternal-fetal interface in rhesus macaques"

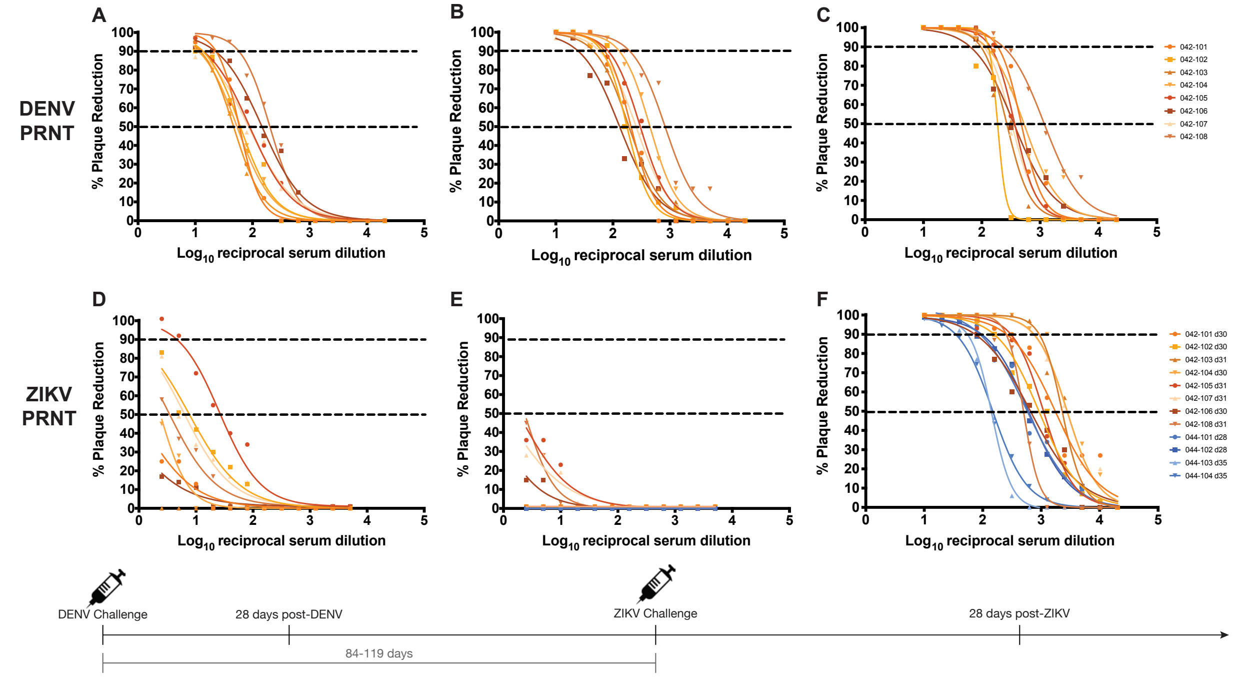


**Fig. S1. PRNT neutralization curves.** PRNT titers against DENV (**A-C**) and ZIKV (**D-E**) at 28 days post-DENV challenge, 0 days post-ZIKV challenge, and 28-35 days post ZIKV challenge.

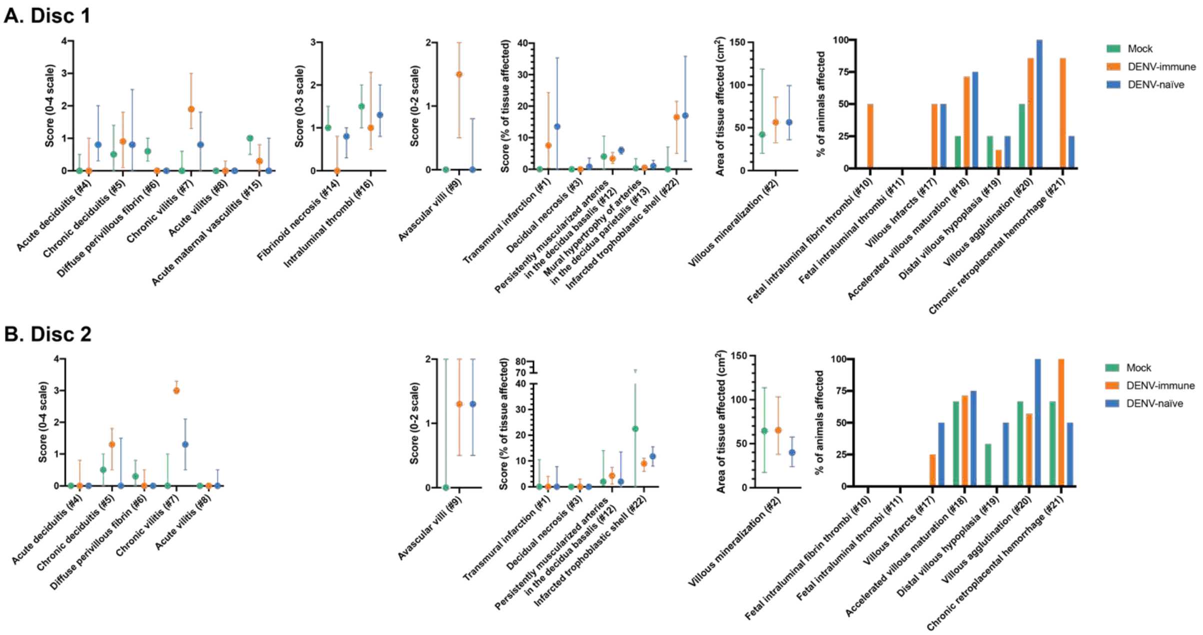


**Fig. S2. Placental pathology scoring.** The central cross-section of each placental disc was evaluated for 22 pathologic changes. A description of the scoring system can be found in Supplementary Table 4.  Features specific to the fetal membranes or uterus are noted in disc 1 scoring. Statistical pairwise comparisons between each group were performed for each feature. For quantitative features (1-9, 12-16, 22) a non-parametric Wilcoxon rank sum test was used; for binary features (10-11, 17-21) Fisher's exact test was used. For quantitative features, the median value is shown with error bars representing the interquartile range. When compared to the mock-infected cohort, the DENV-immune macaques had significantly higher scores for transmural infarction (disc 1: p=0.0371; disc 2: not significant), chronic villitis (disc 1: p=0.0207; disc 2: p=0.0151), avascular villi (disc 1: p=0.0152; disc 2: not significant), and chronic retroplacental hemorrhage (disc 1: p=0.0152; disc 2: not significant).

### Table S1. Maternal and Fetal Tissue and Fluid ZIKV RNA Detection

| | | # positive/total tested | |
|---|---|---|---|
| | **Tissue** | **DENV-immune** | **DENV-naïve** |
| **Maternal** | Mesenteric lymph node | 0/8 | 1/4 |
| | Spleen | 0/2<br>(042-101, 042-103 not collected) | not collected |
| | Liver | 0/2<br>(042-101, 042-103 not collected) | not collected |
| **Fetal** | Umbilical cord plasma | 0/8 | 0/4 |
| | Amniotic fluid | 0/7<br>(042-504 not collected) | 0/4 |

Spleen and liver biopsies were not collected from the majority of dams in order to minimize the size of the c-section incision.

### Table S2. Placental Pathology Scoring System (central section)

| Category | # | Feature | Possible scores |
|---|---|---|---|
| General | 1 | Transmural infarction | 0-100% |
| | 2 | Villous mineralization/cm$^2$ (20x magnification) | area (/cm$^2$) |
| | 3 | Decidual necrosis | 0-100% |
| | 4 | Acute deciduitis | 0-4 |
| | 5 | Chronic deciduitis | 0-4 |
| | 6 | Diffuse perivillous fibrin | 0-4 |
| Villitis | 7 | Chronic villitis: presence of lymphocytes, macrophages | 0-4 |
| | 8 | Acute villitis: presence of neutrophils | 0-4 |
| Fetal vascular malperfusion | 9 | Avascular villi (high grade = 2; low grade = 1; insignificant = 0) | 0-2 |
| | 10 | Fetal intraluminal fibrin thrombi in the stem villlous, chorionic plate vessel, or umbilical cord (present = 1; absent = 0) | 0-1 |
| | 11 | Fetal intramural thrombi in the stem villlous, chorinic plate vessel, or umbilical cord (present = 1; absent = 0) | 0-1 |
| Maternal vascular malperfusion | 12 | Persistently muscularized arteries in the decidua basalis | 0-100% |
| | 13 | Mural hypertrophy of fetal membrane arteries in the decidua parietalis | 0-100% |
| | 14 | Fibrinoid necrosis of vessels in the decidua parietalis, decidua basalis, or trophoblastic shell of the basal plate (present = 1/site; max 3) | 0-3 |
| | 15 | Acute maternal vasculitis in the uterus, decidua parietalis, decidua basalis, or trophoblastic shell of basal plate (present = 1/site; max 4) | 0-4 |
| | 16 | Intraluminal fibrin thrombi in the uterus, decidua parietalis, or decidua basalis (present = 1/site; max 3) | 0-3 |
| | 17 | Villous infarcts (present = 1; absent = 0) | 0-1 |
| | 18 | Accelerated villous maturation (AVM): small villi usually with increased syncytial knots (present = 1; absent = 0) | 0-1 |
| | 19 | Distal villous hypoplasia: villi are the same size as AVM but fewer very small tertiary villi and greater intervillous space indicating chronicity (present = 1; absent = 0) | 0-1 |
| | 20 | Villous agglutination: Maternal vascular malperfusion and/or chronic villitis (present = 1; absent = 0) | 0-1 |
| | 21 | Chronic retroplacental hemorrhage: hemosiderin in basal plate or decidua basalis (present = 1; absent = 0) | 0-1 |
| | 22 | Infarcted trophoblastic shell | 0-100% |
